# Sex-specific additive genetic variances and correlations for fitness in a song sparrow (*Melospiza melodia*) population subject to natural immigration and inbreeding

**DOI:** 10.1101/272138

**Authors:** Matthew E. Wolak, Peter Arcese, Lukas F. Keller, Pirmin Nietlisbach, Jane M. Reid

## Abstract

Quantifying sex-specific additive genetic variance (V_A_) in fitness, and the cross-sex genetic correlation (r_A_), is pre-requisite to predicting evolutionary dynamics and the magnitude of sexual conflict. Quantifying V_A_ and r_A_ in underlying fitness components, and multiple genetic consequences of immigration and resulting gene flow, is required to identify mechanisms that maintain V_A_ in fitness. However, these key parameters have rarely been estimated in wild populations experiencing natural environmental variation and immigration. We used comprehensive pedigree and life-history data from song sparrows (*Melospiza melodia*) to estimate V_A_ and r_A_ in sex-specific fitness and underlying fitness components, and to estimate additive genetic effects of immigrants as well as inbreeding depression. We found substantial V_A_ in female and male fitness, with a moderate positive cross-sex rA. There was also substantial V_A_ in adult reproductive success in males but not females, and moderate V_A_ in juvenile survival but not adult survival. Immigrants introduced alleles for which additive genetic effects on local fitness were negative, potentially reducing population mean fitness through migration load, yet alleviating expression of inbreeding depression. Substantial V_A_ for fitness can consequently be maintained in the wild, and be concordant between the sexes despite marked sex-specific V_A_ in reproductive success.

## Introduction

The magnitude of additive genetic variance (V_A_) in fitness governs the rate of adaptive trait evolution and the expected increase in population mean fitness (Fisher 1930; Robertson 1966; Price 1970), and thereby links adaptation and population persistence (Bell 2013; Gomulkiewicz and Shaw 2013; Carlson et al. 2014; Shaw and Shaw 2014). Quantifying the magnitude of V_A_ in fitness, and identifying key mechanisms that maintain or constrain such V_A_, are consequently central objectives in evolutionary biology (Burt 1995; Barton and Keightley 2002; Ellegren and Sheldon 2008; Walsh and Blows 2009; Shaw and Shaw 2014).

The magnitude and maintenance of V_A_ will partly depend on the genetic architecture underlying fitness in and among individuals. Specifically, in organisms with separate sexes, many genes that affect fitness will be expressed in both sexes. Such genes can have congruent or divergent pleiotropic effects on multiple sex-specific fitness components encompassing survival to sexual maturity and subsequent reproductive success (Arnold and Wade 1984; Falconer 1989, p.338; Chippindale et al. 2001). Such positive or negative pleiotropy can create additive genetic correlations (r_A_) between the sexes, and among fitness components within each sex, potentially generating evolutionary sexual conflict and multiple life-history trade-offs and multi-dimensional constraints (Lande 1980, 1982; Rose 1982; Charlesworth 1987; Chippindale et al. 2001; Kruuk et al. 2008; Bonduriansky and Chenoweth 2009; Walsh and Blows 2009; Shaw and Shaw 2014). Consequently, V_A_ in sex-specific fitness and fitness components, and corresponding cross-sex and within-sex r_A_s, are key parameters that shape the total V_A_ for fitness that emerges and is maintained following selection (Lewontin 1974; Rose 1982; Chippindale et al. 2001; Brommer et al. 2007; Kruuk et al. 2008; Walsh and Blows 2009; Walling et al. 2014).

Further, the magnitude of V_A_ in fitness and underlying fitness components that is maintained in any focal population or sub-population will also depend on natural spatio-temporal variation in fitness, and on resulting variation in the form of local selection and adaptation and associated patterns of immigration and inter-deme gene flow (Merilä and Sheldon 1999; Zhang 2012; Carlson et al. 2014; Shaw and Shaw 2014). Immigration could increase V_A_ by introducing alleles with negative or positive additive effects on local fitness, potentially causing migration load, and impeding or facilitating adaptation and population growth (Lenormand 2002; Garant et al. 2007; Edelaar and Bolnick 2012; Carlson et al. 2014). Such immigration could further change mean fitness by altering the degree of local inbreeding versus outbreeding and associated expression of inbreeding depression, heterosis, and outbreeding depression (Ingvarsson and Whitlock 2000; Tallmon et al. 2004; Frankham 2016). Such effects, and resulting effective rates of gene flow, depend fundamentally on the genetic properties of immigrants relative to focal natives (Ingvarsson and Whitlock 2000; Tallmon et al. 2004; Edelaar and Bolnick 2012). Therefore, understanding and predicting overall evolutionary dynamics requires estimation of V_A_ in fitness and underlying fitness components in both sexes, and associated cross-sex and within-sex rAs, in wild populations experiencing natural abiotic and biotic environmental variation (Ellegren and Sheldon 2008; Kirkpatrick 2009; Kruuk et al. 2008, 2014; Shaw and Shaw 2014; Walling et al. 2014), and also requires explicit estimation of multiple genetic effects resulting from immigration (Ingvarsson and Whitlock 2000; Lenormand 2002; Tallmon et al. 2004; Garant et al. 2007; Edelaar and Bolnick 2012; Carlson et al. 2014).

Fitness can be defined and measured in numerous ways (Brommer 2000; Metcalf and Pavard 2007; Orr 2009; Sæther and Engen 2015). In the context of Fisher’s (1930) Fundamental Theorem, absolute fitness is most straightforwardly defined as the total number of zygotes produced by a zygote (Crow and Kimura 1970; Arnold and Wade 1984; Falconer 1989 p. 336; Shaw and Shaw 2014). Such fitness emerges from sequential life-history events, encompassing survival from conception to sexual maturity and subsequent adult lifetime reproductive success (LRS). In iteroparous species, adult LRS itself results from a repeating sequence of reproduction followed by survival towards the next reproductive opportunity, terminated by death. Fitness and its components therefore reflect expression of numerous developmental, physiological, morphological and behavioral traits, and are consequently best conceptualized as highly polygenic, complex traits (e.g. Houle 1992; Barton and Keightley 2002; Flint and Mackay 2009; Hill 2012; Travisano and Shaw 2013) even though loci of large effect can exist (e.g. Johnston et al. 2013; Trask et al. 2016). Key V_A_s and r_A_s can be estimated using quantitative genetic methods derived from the infinitesimal model (Lynch and Walsh 1998). Although the phenotypic distribution of fitness is intrinsically non-Gaussian (Arnold and Wade 1984; Wagenius et al. 2010; Shaw and Etterson 2012; Bell 2013; Shaw and Shaw 2014), V_A_s and r_A_s can be estimated on appropriate latent scales in order to fulfill the fundamental quantitative genetic assumption of multivariate normality of the average effect of an individual’s polygenic genotype (i.e. breeding value, Lynch and Walsh 1998 pp.72–79; de Villemereuil et al. 2016).

Such estimation of V_A_s and r_A_s in wild populations is empowered by a class of quantitative genetic generalized linear mixed models (QGGLMMs) commonly known as ‘animal models’ (Kruuk 2004; Charmantier et al. 2014). Such models partition variance in observed phenotypes across individuals and, given an appropriately specified relatedness matrix and model, minimize biases in estimates of V_A_ and r_A_ stemming from selection (i.e. non-random variation in fitness) and resulting unobservable phenotypes, as well as estimate variances arising from shared environmental effects (Henderson 1973; Kruuk 2004; Kruuk and Hadfield 2007; Hadfield 2008). Such models can also directly estimate mean additive genetic values of immigrants relative to natives, and estimate the magnitude of inbreeding depression, and thereby elucidate key roles of immigration and resulting gene flow in shaping phenotypic means and variances (Reid and Keller 2010; Wolak and Keller 2014; Wolak and Reid 2017).

However, despite the widely recognized need and available statistical methods, few studies have rigorously estimated sex-specific V_A_s and the cross-sex r_A_ in fitness in wild populations (Burt 1995; Gardner et al. 2005; Kruuk et al. 2008; Kirkpatrick 2009; Shaw and Shaw 2014). Of 17 known studies (including on humans, *Homo sapiens*) that estimated V_A_ for sex-specific absolute fitness measured approximately zygote to zygote, only 8 considered male fitness alongside female fitness (Appendix S1). Since most such studies estimated at least one sex-specific V_A_ to be close to zero, possibly due to low power, only two attempted to estimate the cross-sex r_A_ (McFarlane et al. 2014; Zietsch et al. 2014, Appendix S1). Two further studies estimated the cross-sex r_A_ for fitness measured as an adult’s number of adult (i.e. recruited) offspring (Brommer et al. 2007; Foerster et al. 2007), but such cross-generation measures are harder to reconcile with primary evolutionary theory (Arnold and Wade 1984; Wolf and Wade 2001), and all available estimates are very imprecise (Appendix S1). Further, few studies explicitly estimated V_A_ on appropriate latent scales (but see Milot et al. 2011; McFarlane et al. 2014) or then transformed estimates back onto observed phenotypic scales, as is ideally required to facilitate cross-study comparison (de Villemereuil et al. 2016). Finally, no QGGLMM analysis of V_A_ in fitness has explicitly estimated additive genetic effects of immigrants, or thereby directly assessed the role of introgressive gene flow in changing local mean breeding value and maintaining V_A_ and associated evolutionary potential (Wolak and Reid 2016, 2017).

This paucity of estimates likely reflects the substantial challenges of collecting comprehensive sex-specific fitness and relatedness data from free-living individuals. Since wild population studies can rarely count all conceived zygotes, fitness can be pragmatically quantified as the total number of offspring produced over an individual’s lifetime, where focal individuals and their offspring are censused as close to conception as feasible (typically soon after birth, hatch, or seed formation). However, most field datasets have some degree of missing or incorrect parentage assignment, and resulting pedigree error could bias quantitative genetic analyses (Brommer et al. 2007; Firth et al. 2015; Wolak and Reid 2017). Further, challenges of tracking juveniles among natal and subsequent breeding locations, and of paternity assignment, mean that records of survival to maturity and male reproductive success are often missing or incorrect (Kruuk et al. 2000; Brommer et al. 2007; Stinchcombe 2014). Observed distributions of fitness may also exclude unobserved non-breeders, and hence inaccurately reflect the frequency of individuals with zero fitness. Such error will likely bias, with respect to fitness, estimates of VA, phenotypic means and variances, and hence key standardized metrics that depend on V_A_ and moments of the phenotypic distribution and that underpin comparative analyses (heritability, h^2^; evolvability, I_A_; coefficient of additive genetic variance, CV_A_; e.g. Freeman-Gallant et al. 2005). Even given comprehensive data spanning multiple generations, V_A_s and cross-sex r_A_s in non-Gaussian traits are notoriously difficult to estimate precisely (Shaw 1987; Poissant et al. 2010; Kruuk et al. 2014). Statistical methods that adequately quantify uncertainty should then be used to facilitate inference and subsequent meta-analyses (Garcia-Gonzalez et al. 2012).

Accordingly, we fitted Bayesian QGGLMMs to comprehensive multi-generation fitness and pedigree data from song sparrows (*Melospiza melodia*) to estimate sex-specific V_A_s and r_A_s in fitness, and in two hierarchical levels of fitness components. Specifically, we estimated (*i*) V_A_ in sex-specific fitness and the cross-sex r_A_, thereby evaluating scope for inter-sexual conflict; (*ii*) V_A_ and r_A_ in and among juvenile survival and sex-specific adult LRS, comprising the primary fitness components that generate overall fitness; and (*iii*) V_A_ and r_A_ in sex-specific adult annual reproductive success (ARS) and (*iv*) V_A_ in adult annual survival, comprising the key life-history traits that generate adult LRS. In all cases, we explicitly estimated additive genetic effects of immigrants relative to defined local population founders, and estimated the magnitude of inbreeding depression, and thereby evaluate concurrent impacts of natural immigration and resulting gene flow on local additive genetic and phenotypic variation in fitness.

## Materials and Methods

### STUDY SYSTEM

Estimating V_A_ and r_A_ in fitness and fitness components in the wild is perhaps most tractable in populations with limited emigration but sufficient immigration to generate substantial variance in relatedness, and where all local residents and immigrants can be observed. A population of song sparrows inhabiting Mandarte Island, British Columbia, Canada, fulfills these criteria and has proved valuable for quantifying fitness of residents and immigrants and for pedigree-based quantitative genetic analyses (Keller 1998; Marr et al. 2002; Reid et al. 2011, 2014a,b; Reid and Sardell 2012; Wolak and Reid 2016).

Mandarte’s song sparrows typically form socially monogamous breeding pairs, starting from age one year, with a mean of 28±11SD (range 11–52) breeding females per year during 1993–2015. Pairs can rear up to three broods of chicks per year (mean brood size 2.8±1.0SD chicks, range 1–4). However, 28% of offspring are sired by extra-pair males (Sardell et al. 2010), creating opportunities for individual males to gain or lose substantial reproductive success compared to their socially-paired female (Reid et al. 2011, 2014b; Reid and Sardell 2012). Further, since the adult sex-ratio is often male-biased (mean proportion males during 1993–2015: 0.60±0.09SD, range 0.39–0.75), some males remain socially unpaired in some years (Lebigre et al. 2012), and these males typically gain little extra-pair paternity (Sardell et al. 2010). Consequently, the population’s mating system and ecology fosters different means and variances in female versus male reproductive success (Lebigre et al. 2012), creating potential for sexual conflict and trade-offs over fitness components despite social monogamy.

Since 1975, virtually all song sparrow breeding attempts on Mandarte were closely monitored and all chicks surviving to ca. 6 days post-hatch were marked with unique combinations of metal and colored plastic bands (Smith et al. 2006). Mandarte lies within a large song sparrow metapopulation and receives occasional immigrants (totaling 28 females and 16 males during 1976–2014) that were mist-netted and color-banded soon after arriving (Marr et al. 2002; Reid et al. 2006; Smith et al. 2006). Consequently, every song sparrow in the population is individually identifiable by field observation. Comprehensive surveys undertaken each April identified all surviving individuals, including unpaired males, with resighting probability >0.99 (Wilson et al. 2007). Local chick survival from banding to adulthood the following April, and adult survival to subsequent years, were consequently accurately recorded (Keller 1998; Smith et al. 2006).

Each year, the socially-paired parents that reared all banded offspring were identified. To determine genetic parentage, since 1993 all banded chicks and adults were blood sampled and genotyped at 160 polymorphic microsatellite loci. All chicks were assigned to genetic parents with >99% individual-level confidence (Sardell et al. 2010; Nietlisbach et al. 2015). These analyses demonstrated zero extra-pair maternity, and effectively eliminated paternity error. Each banded individual’s sex was determined from adult reproductive behavior and/or by genotyping the chromobox-helicase-DNA-binding (CHD) gene (Postma et al. 2011; Nietlisbach et al. 2015).

The local fitness of each chick banded on Mandarte since 1993 was measured as its total lifetime number of chicks banded on Mandarte, including zeros for chicks that died before adulthood (Appendix S2). The two major fitness components, juvenile survival and adult LRS, were respectively measured as survival from banding to adulthood the following April, and the total number of banded chicks assigned to individuals that survived to adulthood. For each adult, LRS was then further subdivided into ARS and annual survival, respectively measured as the number of banded chicks assigned to each individual in any one year, and survival to the following April. Since adult (breeding) dispersal away from Mandarte is probably very rare, observed local adult survival likely equates to true survival (Marr et al. 2002; Smith et al. 2006). The relatively high local recruitment rate implies that juvenile (natal) dispersal is also relatively infrequent, although non-zero. However, surveys of immediately surrounding islands have detected few local dispersers, implying that unobserved dispersal from Mandarte is likely to be longer distance. Observed juvenile survival on Mandarte is therefore an appropriate measure of effective local survival and hence local fitness.

### QUANTITATIVE GENETIC MODELS

We fitted a series of four non-Gaussian QGGLMMs designed to estimate sex-specific additive genetic variances (V_A_) and covariances (COV_A_), and hence estimate associated standardized statistics (r_A_, h^2^, I_A_, CV_A_), for fitness and fitness components.

First, we fitted a bivariate QGGLMM (Appendix S4) to estimate V_A_ in female and male fitness and the cross-sex COV_A_, assuming Poisson distributions with log link functions. Random hatch-year effects were fitted to estimate sex-specific cohort variances in fitness and the cross-sex cohort covariance. Sex-specific residual variances were estimated assuming additive overdispersion, with residual covariance fixed to zero.

Second, we fitted a trivariate QGGLMM (Appendix S4) to estimate V_A_ in juvenile survival and adult female and male LRS, and the three pairwise COV_A_s. We modeled juvenile survival as a single joint trait of both sexes with sex-specific intercepts, rather than as two sex-specific traits. This simplification facilitated multivariate analysis of juvenile survival alongside sex-specific adult LRS, and is justified because previously published and exploratory analyses demonstrated a strong positive cross-sex r_A_ for juvenile survival and similar magnitudes of V_A_ in both sexes, implying considerable shared V_A_ (Reid and Sardell 2012, Appendix S7). Under these conditions, modeling a single trait for both sexes does not bias estimates of V_A_ (Wolak et al. 2015). Juvenile survival was modeled as a binary trait with logit link function and residual variance fixed to one. We assumed Poisson distributions for female and male LRS, with log link functions and independent residual variances (as for fitness). Random hatch-year effects were again fitted, thereby estimating cohort variances and covariances in and among the three traits. Random effects of the identities of each chick’s mother, social father, social parent pair, and brood were also fitted for juvenile survival, thereby accounting for common environmental effects stemming from parental care and natal conditions. While these four effects may be somewhat confounded, our aim was not to precisely estimate associated variances, but simply to minimize possible bias in V_A_ (e.g. Kruuk and Hadfield 2007; Reid et al. 2014a). Analogous common environmental effects were not fitted to female and male adult LRS, because relatively few parents and broods produced multiple same-sex chicks that survived to adulthood, and previous analyses did not reveal substantial parental effects on adult life-history traits.

Third, we fitted a bivariate QGGLMM (Appendix S4) to estimate V_A_ in adult female and male ARS and the cross-sex COV_A_ assuming Poisson distributions for both traits, log link functions, and independent residual variances. Random individual effects were fitted to estimate sex-specific permanent individual variances (i.e. repeatable among-individual variation stemming from permanent environmental and/or non-additive genetic effects). Random year of observation effects were also fitted to estimate among-year environmental variances and the cross-sex year covariance.

Fourth, we fitted a univariate QGGLMM (Appendix S4) to estimate V_A_ in annual adult survival modeled as a single trait for both sexes with sex-specific intercepts (as for juvenile survival). We modeled survival as a binary trait expressed by each individual adult in each year, with logit link function and residual variance fixed to one (e.g. Hadfield et al. 2013). Random year of observation and individual effects were fitted to estimate among-year environmental variance and account for overdispersion compared to the assumed geometric distribution of age-specific survival events.

### IMMIGRANTS, INBREEDING DEPRESSION, AND FIXED EFFECTS

Standard QGGLMMs estimate V_A_ and COV_A_ for a default base population that comprises ‘phantom parents’ of all pedigreed individuals with unknown parents (Kruuk 2004; Wolak and Reid 2017). In populations with complete local pedigree data for a focal study period but that are open to immigration, the default base population comprises phantom parents of all adults alive at the study start (hereafter ‘founders’) and of subsequent immigrants. To directly estimate the difference in mean additive genetic value for fitness and fitness components between the defined founders and subsequent immigrants, and account for heterogeneity that could otherwise bias V_A_ estimates, all four QGGLMMs included trait-specific linear regressions on individual immigrant genetic group (*IGG*) coefficient. Each individual’s *IGG* coefficient quantifies the expected proportion of that individual’s autosomal genome that originated from the defined immigrant group, calculated from pedigree data (Appendix S3). The regression slope (β*_IGG_*), modeled as a fixed effect, estimates the difference in mean additive genetic value of the immigrant group relative to the founder group (Wolak and Reid 2017). Since immigration was infrequent, phantom parents of female and male immigrants that arrived in all years were pooled into a single genetic group (Appendix S3). This assumes that the phantom mothers of female and male immigrants have similar mean genetic values as the phantom fathers for any focal trait, and hence that alleles originating in immigrants of both sexes similarly affect the genetic values of descendants of both sexes. This mirrors the standard QGGLMM assumption that female and male phantom parents of founders have the same mean breeding values for any focal trait (Wolak et al. 2015).

To quantify inbreeding depression, and minimize bias in V_A_ estimates that can result from correlated inbreeding across relatives, all four QGGLMMs also included trait-specific linear regressions on individual coefficient of inbreeding (*f*), calculated from pedigree data (Reid and Keller 2010; Wolak and Keller 2014). Regression slopes (β*_f_*) equate to haploid ‘lethal equivalents’ for traits modeled with log link functions (fitness, adult LRS, and ARS), but not for traits modeled with logit link functions (juvenile and adult survival).

Further fixed effects were restricted to those required to standardize trait observations across individuals. Since juvenile survival probability decreases with increasing seasonal hatch date (Smith et al. 2006), and hatch date reflects the parents’ breeding phenotype, models for juvenile survival included a linear regression on the first egg lay date in the nest in which each focal individual hatched. Since adult ARS and survival vary with age (Smith et al. 2006; Keller et al. 2008), associated models included categorical effects of age at observation (ages 1, 2, 3–5, or ≥6 years).

### PEDIGREE DATA AND MODEL IMPLEMENTATION

Comprehensive pedigree data were initially compiled by assigning all offspring banded during 1975–2014 to their observed socially-paired parents. Paternal links for all chicks hatched during 1993–2014, and 37 additional genotyped chicks hatched during 1991–1992, were then corrected for extra-pair paternity (Sardell et al. 2010; Reid et al. 2011; Nietlisbach et al. 2015, 2017). For each QGGLMM, the pedigree was pruned to individuals with observed phenotypes and their known ancestors. The inverse numerator relationship matrix, and individuals’ *IGG* and *f* coefficients, were computed using standard algorithms (Wolak and Reid 2017, Appendix S3). Immigrants were defined as unrelated to all Mandarte residents at arrival, and to subsequent immigrants (Marr et al. 2002; Reid et al. 2006).

For each model, phenotypic data were restricted to cohorts for which all or virtually all individuals had complete fitness or fitness component data, known sex, and genetically verified parents (Appendix S2). Observations of immigrants’ own phenotypes were excluded because they might reflect ecological effects associated with dispersal or subsequent settlement (Marr et al. 2002), and because immigrants’ *f* values are undefined relative to the Mandarte pedigree base population (Reid et al. 2006). However, immigrants that produced ≥1 banded offspring were explicitly included in the pedigree to enable estimation of relatedness among descendants and genetic group effects.

All models were implemented in a Bayesian framework, using a Markov chain Monte Carlo (MCMC) algorithm to sample posterior distributions. We used diffuse normal prior distributions for all fixed effects (mean=0, variance=10^10^), and multivariate parameter expanded priors for covariance matrices that gave uniform marginal prior distributions on the correlation. Parameter expanded priors were used for other variance components, giving scaled non-central F-distributions with numerator and denominator degrees of freedom of one (Gelman 2006; Hadfield 2010) and scale parameter of 10 for binary traits or 1,000 for Poisson traits (Appendix S4).

We retained 5,000 samples of each marginal posterior distribution, with MCMC burn-in and thinning interval set to yield absolute autocorrelation values <0.1 and satisfy convergence criteria (Appendix S4). When marginal posterior distributions are approximately Gaussian, posterior modes and 95% highest posterior density credible intervals (95%CI) sufficiently summarize point estimates and uncertainty. However, distributions can show skew, kurtosis or multiple peaks, including when parameters are near their boundary (e.g. variance near zero). Inferences drawn from posterior modes versus means may then differ. Consequently, we report the marginal posterior mean, mode, and 95%CI and, for key metrics, depict full marginal posterior distributions alongside prior distributions to facilitate interpretation (Appendix S4).

All QGGLMMs assumed Poisson or binary distributions and therefore estimated (co)variances on latent scales. Posterior distributions of latent-scale heritability 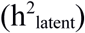 and r_A_ were computed from all samples of the marginal posterior distributions of underlying components following standard formulae (Appendix S4). However, latent-scale statistics are model specific and not directly comparable among analyses or populations, or interpretable on the scale on which phenotypes are expressed and experience natural selection (de Villemereuil et al. 2016). Therefore, to facilitate future comparative studies and evolutionary inferences, we attempted to back-transform posterior distributions of latent-scale variances to the observed phenotypic scale and calculate observed-scale posterior distributions of standardized summary statistics (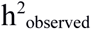, I_A-observed_, CV_A-observed_, Appendices S4, S5). However, we could not recover reliable observed-scale variance component posteriors from our first model of female and male fitness due to the substantial overdispersion (Appendix S2). I_A-observed_ was not calculated for juvenile or adult survival because mean standardized variances are not meaningful for binary traits where the mean phenotype is bounded by 0 and 1 (Houle 1992).

Analyses were conducted in R (v3.2.3, R Core Team 2015) using the MCMCglmm (v2.22.1, Hadfield 2010), nadiv (v2.14.3.2, Wolak 2012) and QGglmm (v0.6.0, de Villemereuil et al. 2016) packages. Additional univariate QGGLMMs for sex-specific fitness, and univariate and bivariate QGGLMMs for combinations of juvenile survival and adult LRS, gave quantitatively similar variance component estimates as the four main QGGLMMs. Key (co)variance component estimates are robust to reasonable alternative priors (Appendix S6), and remained similar when additional parental environmental effects were modeled. Additional details of results, and descriptive figures, are in Appendix S5. Data and R code will be available on Dryad and GitHub (https://github.com/matthewwolak/Wolak_etal_SongSparrowFitnessQG), once the manuscript is accepted for publication.

## Results

### FITNESS

Across 1406 female and 1415 male chicks banded on Mandarte during 1993–2012, 1177 (83.7%) and 1185 (83.7%) respectively had zero fitness. Consequently, fitness distributions were strongly right-skewed, with maxima of 50 and 69 banded offspring for females and males respectively (Fig. 1A). Raw mean sex-specific fitness was 1.78 and 1.70 respectively, with substantial phenotypic variances (females 29.8, males 31.7).

**Figure 1.**
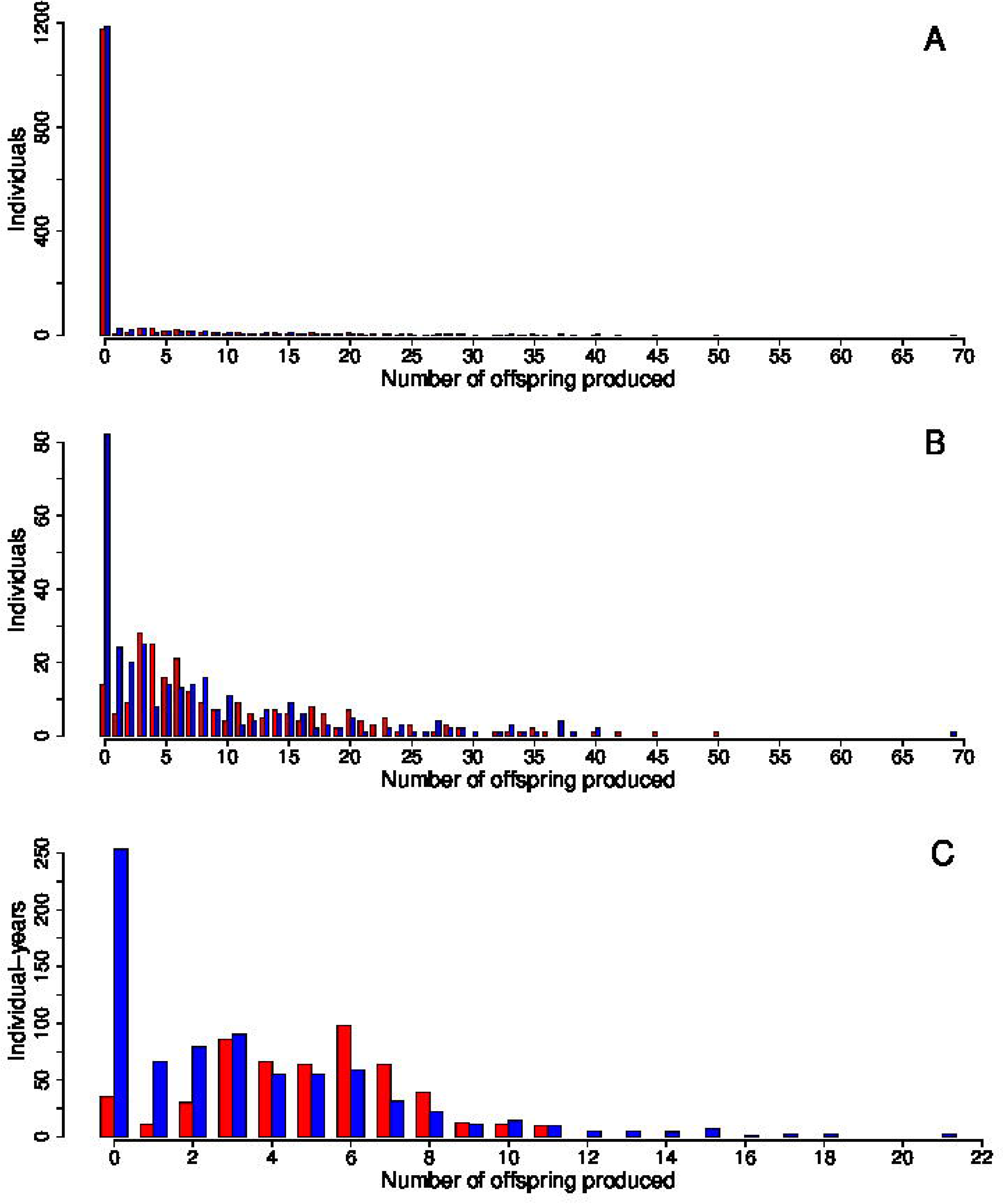
Phenotypic distributions of (A) fitness, (B) adult lifetime reproductive success, and (C) adult annual reproductive success measured as the number of banded chicks attributed to each focal individual. Red and blue denote females and males respectively.

In the bivariate QGGLMM, the posterior distributions for latent-scale V_A_ in female and male fitness showed clear peaks that were substantially shifted away from zero and from the prior distributions, indicating substantial V_A_ for sex-specific fitness (Figs. 2A,B). The posterior modes were similar in both sexes, and the lower 95%CI limits did not converge towards zero (Table 1). There was non-zero cohort variance and substantial residual variance in both sexes, reflecting the overdispersed phenotypic distributions (Table 1, Fig. 1). Consequently, there was relatively small but non-zero heritability of fitness in both sexes; posterior modes and means for 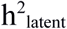 were 0.08–0.09, with lower 95%CI limits that did not converge to zero (Table 1, Fig. S2).

**Figure 2.**
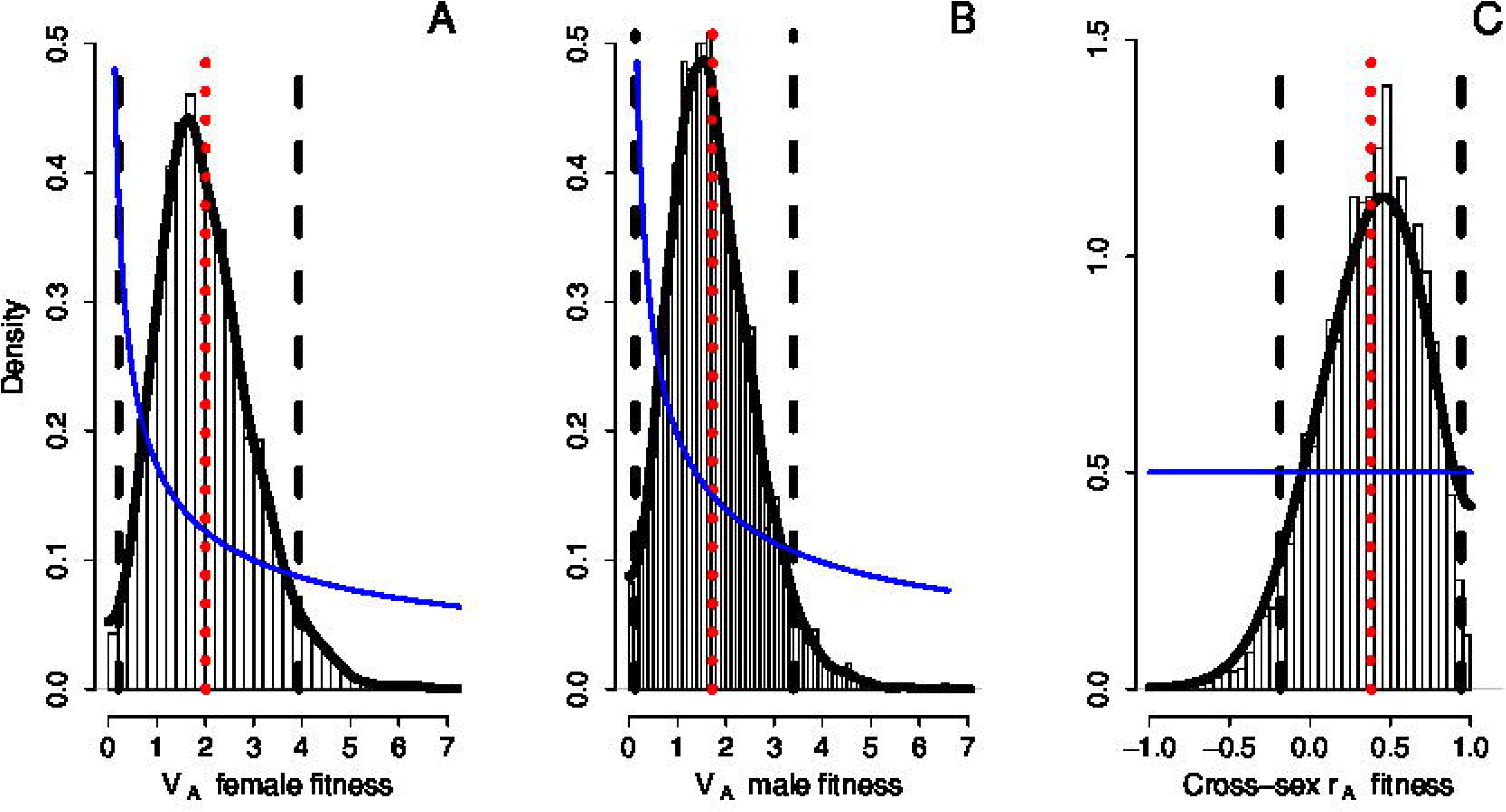
Marginal posterior MCMC samples (bars), kernel density estimation (solid black line), posterior mean (red dotted line), 95% credible interval limits (black dashed lines), and prior (solid blue line) for the additive genetic variances (V_A_) in (A) female fitness, (B) male fitness, and (C) the cross-sex additive genetic correlation (r_A_) in song sparrows. In A and B, the priors are depicted over the range of each posterior distribution, but extend to substantial positive values.

**Table 1.** Marginal posterior means, modes (in square brackets), and 95% credible intervals (in parentheses) for latent-scale estimates from the bivariate model for female and male fitness. Within the additive genetic and cohort matrices, sex-specific variances are shown along the diagonal (bold) with cross-sex covariances (COV) and correlations (r, italics) above and below the diagonal respectively. Sex-specific residual variances (V_R_), heritabilities 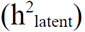, and slopes of regressions on individual coefficient of inbreeding (*β_f_*) and immigrant genetic group coefficient (*β_IGG_*) are also shown.

| | Additive genetic matrix | | Cohort matrix | | $V_R$ | $h^2_{\text{latent}}$ | $\beta_f$ | $\beta_{IGG}$ |
| --- | --- | --- | --- | --- | --- | --- | --- | --- |
|  | Female fitness | Male fitness | Female fitness | Male fitness |  |  |  |  |
| Female<br>fitness | $V_A=2.01$ | $COV_A=0.62$ | **3.12** | $COV=1.46$ | 16.58 | 0.09 | -21.41 | -6.27 |
|  | **[1.56]** | [0.42] | **[2.46]** | [1.16] | [15.29] | [0.08] | [-19.91] | [-5.79] |
|  | **(0.21, 3.93)** | (-0.43, 1.82) | **(0.90, 5.98)** | (0.12, 2.97) | (12.99, 21.06) | (0.02, 0.18) | (-32.53, -11.48) | (-11.36, -2.00) |
| Male<br>fitness | $r_A=0.38$ | $V_A=1.72$ | $r=0.67$ | **1.54** | 15.61 | 0.09 | -27.86 | -5.47 |
|  | [0.45] | **[1.70]** | [0.79] | **[0.97]** | [15.60] | [0.08] | [-25.53] | [-5.19] |
|  | (-0.19, 0.94) | **(0.13, 3.39)** | (0.27, 0.98) | **(0.38, 3.17)** | (11.86, 19.31) | (0.01, 0.17) | (-39.47, -16.50) | (-9.91, -0.71) |

The posterior mode for the cross-sex COV_A_ in fitness was positive, generating a posterior mode for the cross-sex r_A_ of intermediate magnitude between zero and one (Table 1, Fig. 2C). The 95%CI for r_A_ was wide and included zero. However, 88% of the posterior density exceeded zero, representing substantial divergence from the uniform prior, yet the upper 95%CI limit did not converge towards one (Table 1, Fig. 2C). This implies that fitness variation has some, but not all, of the same additive genetic basis in females and males.

In total, 26 immigrants that arrived on Mandarte during 1976–2012 made a non-zero expected genetic contribution to the 2821 Mandarte-hatched individuals whose fitness was observed (Appendix S3). Across these 2821 individuals, mean *IGG* coefficient was 0.52±0.13SD (range 0.14–0.86). Approximately half the focal individuals’ genomes are therefore expected to have originated from immigrants on average, implying that immigration could contribute substantially to standing V_A_ within the Mandarte breeding population. The posterior modes for the regressions of sex-specific fitness on *IGG*, which quantify mean immigrant genetic group effects, were negative in both sexes with 95%CIs that did not overlap zero (Table 1). Additive effects of alleles carried by immigrants therefore decreased fitness, relative to additive effects of alleles in the defined founder population, in both sexes.

Across the 2821 individuals, mean *f* was 0.074±0.052 (range 0.000–0.347, 7.4% zeros). Substantial variation in *f* was directly attributable to immigration: 91% of individuals with *f=0* had one immigrant parent. However, since immigrants’ descendants commonly inbred in future generations, *f* and *IGG* were only moderately correlated across individuals (females: r=-0.25, males: r=-0.30). The posterior modes for the regressions of sex-specific fitness on f were negative with 95%CI that did not overlap zero, demonstrating very strong inbreeding depression in fitness in both sexes (Table 1).

### JUVENILE SURVIVAL AND ADULT LIFETIME REPRODUCTIVE SUCCESS

Of 1542 female and 1562 male chicks banded during 1993–2014, 254 (16.5%) females and 331 (21.2%) males survived on Mandarte to the following April. Adult LRS was measured for 243 adult females and 312 adult males hatched during 1993–2012, with sex-specific means of 10.3 (median 7, variance 85.1, 5.8% zeroes) and 7.7 (median 4, variance 97.6, 26.3% zeroes) banded offspring respectively (Fig. 1B).

In the trivariate QGGLMM, the posterior distribution for V_A_ in juvenile survival showed a clear peak, and hence posterior mean, that departed from zero and from the prior distribution, although the lower 95%CI limit converged towards zero (Table 2, Fig. 3A). There was substantial cohort variance (Table 2), but small variances attributable to mothers, social fathers, broods and parent pairs (Appendix S5). Consequently, the posterior means for 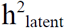 and 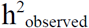 were small, but again showed clear peaks away from zero (Fig. S3). Although the lower 95%CI limits converged towards zero, approximately 93% and 82% of posterior samples respectively exceeded a minimal value of 0.01 (Table 2, Fig. S3).

**Table 2.**
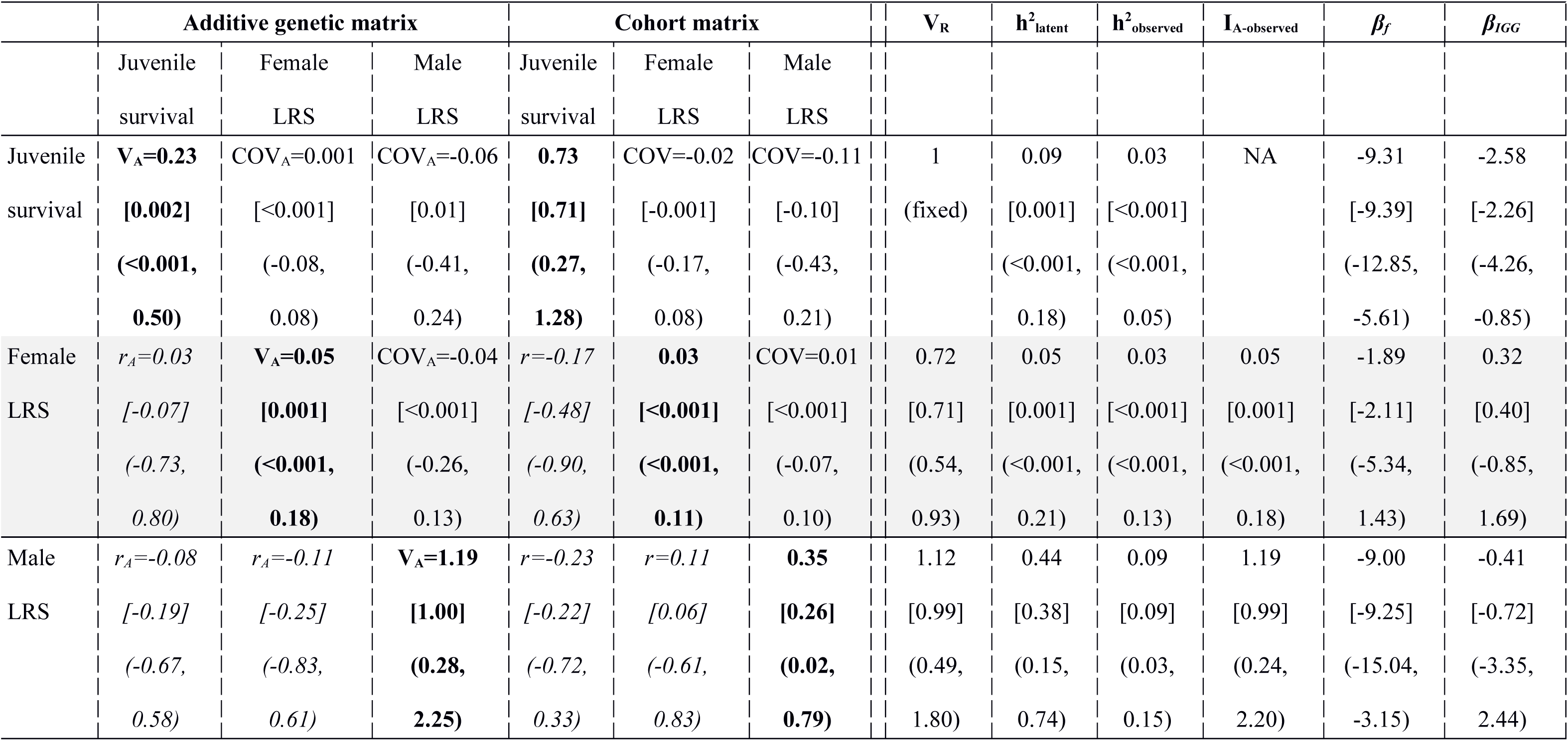
Marginal posterior means, modes (in square brackets), and 95% credible intervals (in parentheses) for latent- and observed-scale estimates from the trivariate model for juvenile survival and adult female and male lifetime reproductive success (LRS). Within the additive genetic and cohort matrices, variances are shown along the diagonal (bold) with covariances (COV) and correlations (r, italics) above and below the diagonal respectively. Residual variances (V_R_), latentscale heritabilities 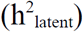, observed-scale heritabilities 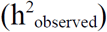 and evolvabilities (I_A-observed_), and slopes of regressions on individual coefficient of inbreeding (*β_f_*) and immigrant genetic group coefficient (*β_IGG_*) are also shown. I_A-observed_ is not applicable (NA) for juvenile survival. Posterior modes and lower 95%CI limits that converged towards zero are reported as <0.001.

**Figure 3.**
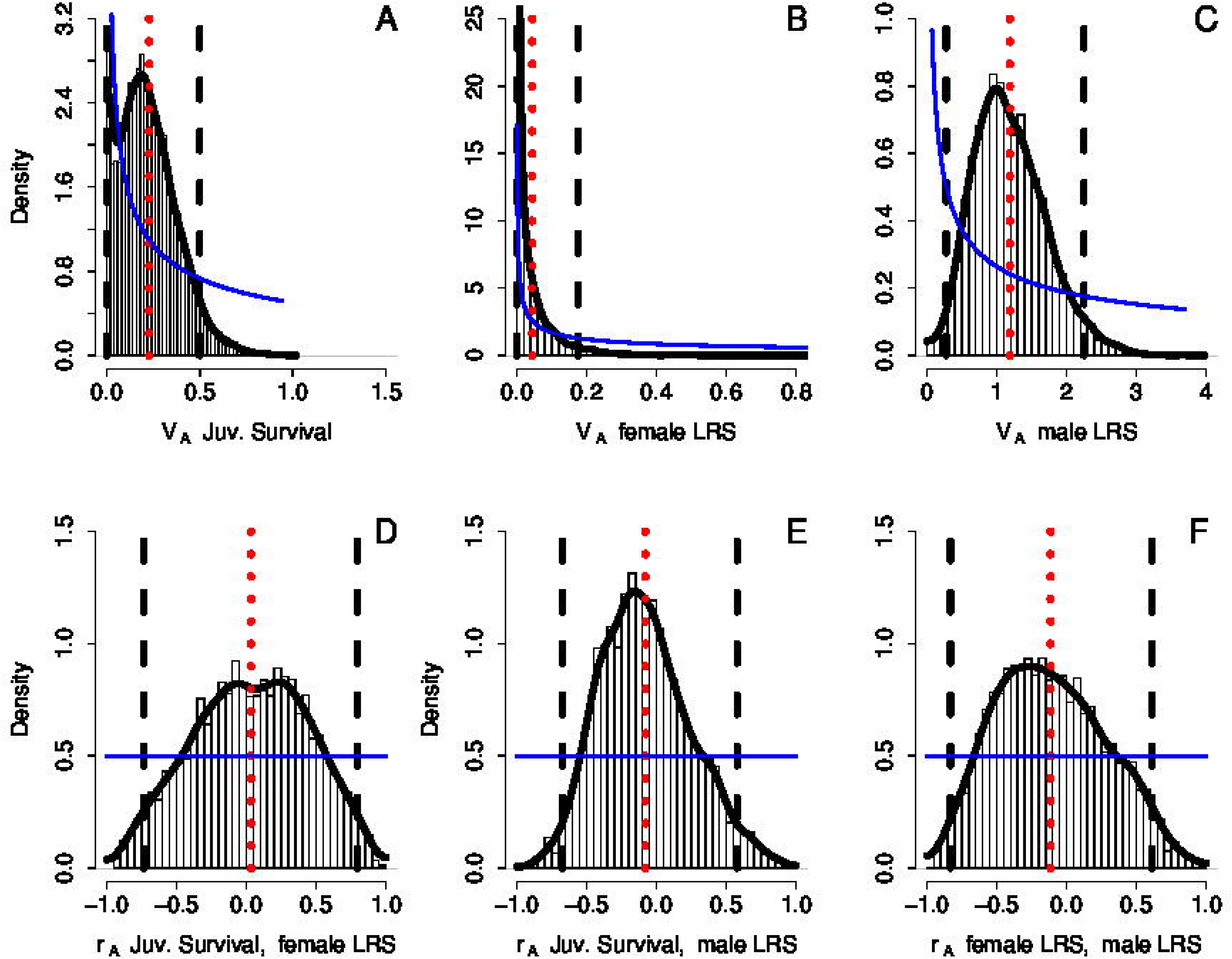
Marginal posterior MCMC samples (bars), kernel density estimation (solid black line), posterior mean (red dotted line), 95% credible interval limits (black dashed lines), and prior (solid blue line) for the additive genetic variances (V_A_) in (A) juvenile survival, (B) adult female lifetime reproductive success (LRS), and (C) adult male LRS, and the additive genetic correlations (r_A_) between (D) juvenile survival and adult female LRS, (E) juvenile survival and adult male LRS, and (F) adult female and male LRS in song sparrows. Note that axis scales vary among plots. In A-C, the priors are depicted over the range of each posterior distribution, but extend to substantial positive values.

The posterior mode for V_A_ in adult female LRS was very small (Table 2). The posterior mean was slightly greater due to the right-skewed posterior distribution (Table 2, Fig. 3B). However there was substantial posterior density close to zero compared to the prior distribution, and the lower 95%CI limit converged towards zero (Fig. 3B, Table 2). Consequently, the posterior modes (and means) of 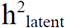, 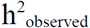 and I_A-observed_ for female LRS were small, with lower 95%CI limits that converged towards zero (Table 2, Figs. S4, S5).

In marked contrast, the posterior mode and mean for V_A_ in adult male LRS were substantial and the lower 95%CI limit considerably exceeded zero (Table 2, Fig. 3C). Consequently, although there were also moderate cohort and residual variances, the posterior mode and mean for 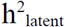 for male LRS were substantial (Table 2, Fig. S4). These values were smaller for 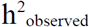, reflecting the non-linear transformation induced by the mean-variance relationship of the Poisson distribution, but the lower 95%CI limit still did not converged towards zero (Table 2, Fig. S4). The posterior mode for I_A-observed_ for male LRS was also moderate (Table 2, Fig. S5).

Since V_A_ in female LRS was so small and the lower 95%CI limit for V_A_ in juvenile survival also converged towards zero, the pairwise COV_A_s and r_A_s among juvenile survival and female and male LRS were unsurprisingly estimated with considerable uncertainty (Table 2, Fig. 3). The posterior modes and means for r_A_ between juvenile survival and male LRS, and between female and male LRS, were slightly negative, but spanned zero for juvenile survival and female LRS, all with 95%CI limits that did not converge towards either -1 or 1 (Table 2, Fig. 3).

Distributions of *IGG* and *f* for individuals included in analyses of juvenile survival and adult LRS (and ARS and survival) were quantitatively similar to those for individuals included in analyses of fitness (summarized above). The posterior mode for the regression of juvenile survival on *IGG* was negative, with a 95%CI that did not overlap zero (Table 2). Further analyses showed similar negative slopes for female and male juvenile survival modeled as separate traits (Appendix S7). However, the posterior modes for the regressions of adult female and male LRS on *IGG* were small, with 95%CIs that spanned zero (Table 2). This implies that additive effects of immigrants’ alleles decreased local juvenile survival, but not adult LRS, relative to additive effects of founders’ alleles.

The posterior modes for the regressions of juvenile survival and adult female and male LRS on *f* were all negative, demonstrating inbreeding depression (although the 95%CI for female LRS overlapped zero, Table 2).

### ADULT ANNUAL REPRODUCTIVE SUCCESS

During 1994–2015, there were 526 and 773 observations of ARS for adult females and males respectively, involving 254 and 331 Mandarte-hatched individuals. Mean female ARS was 4.9 banded offspring (median 5, variance 6.2, range 0–11, 6.7% zeroes, Fig. 1C) and mean male ARS was 3.2 banded offspring (median 2, variance 13.2, range 0–21, 32.7% zeroes, Fig. 1C).

In the bivariate QGGLMM, the posterior mode for V_A_ in female ARS was very small and the lower 95%CI limit converged towards zero (Table 3, Fig. 4A). However, the posterior mean was slightly larger (Table 3), and 75% of the posterior density exceeded a minimal value of 0.01. This implies the existence of very small, but non-zero, V_A_ for female ARS (Fig. 4A inset).

**Table 3.**
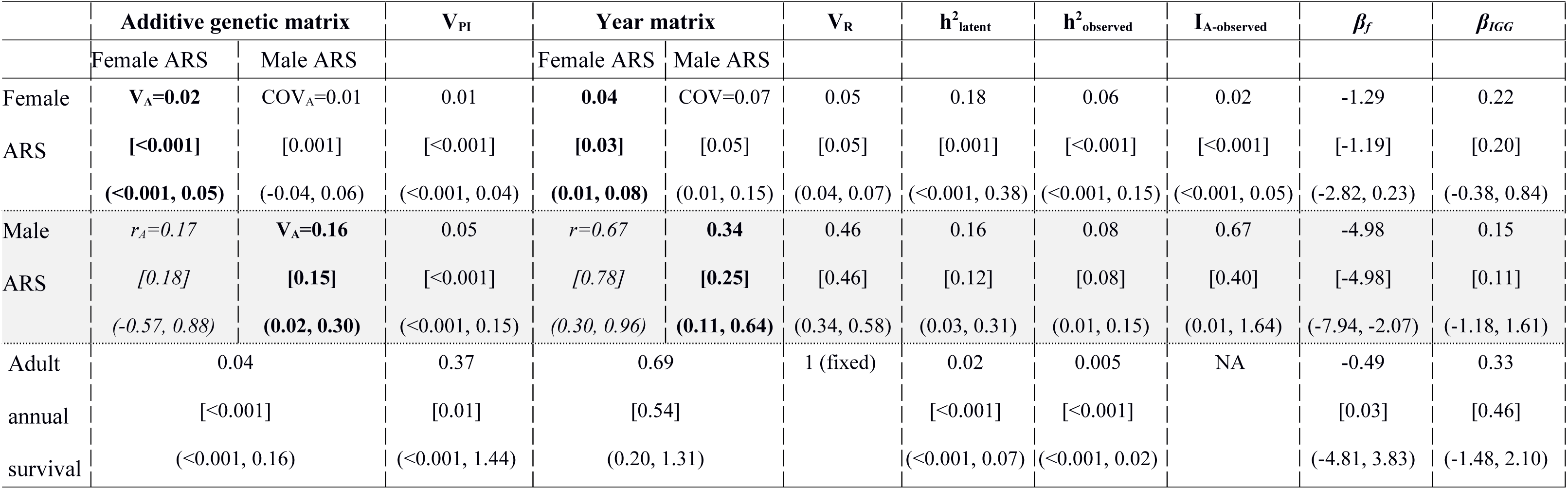
Marginal posterior means, modes (in square brackets), and 95% credible intervals (in parentheses) from (A) the bivariate model for adult female and male annual reproductive success (ARS) and (B) the univariate model for adult annual survival. Within the additive genetic and year matrices for ARS, variances are shown along the diagonal (bold) with covariances (COV) and correlations (r, italics) above and below the diagonal respectively. Permanent individual (V_PI_) and residual (V_R_) variances, latent-scale heritabilities 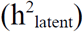, observed-scale heritabilities 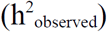 and evolvabilities (I_A-observed_), and slopes of regressions on individual coefficient of inbreeding (β*_f_*) and immigrant genetic group coefficient (*β_IGG_*) are also shown. I_A_ is not applicable (NA) for adult survival. Posterior modes and lower 95%CI limits that converged towards zero are reported as <0.001.

| | Additive genetic matrix | | $V_{PI}$ | Year matrix | | $V_R$ | $h^2_{\text{latent}}$ | $h^2_{\text{observed}}$ | $I_{A\text{-observed}}$ | $\beta_f$ | $\beta_{IGG}$ |
| --- | --- | --- | --- | --- | --- | --- | --- | --- | --- | --- | --- |
|  | Female ARS | Male ARS |  | Female ARS | Male ARS |  |  |  |  |  |  |
| Female | $V_A=0.02$ | $COV_A=0.01$ | 0.01 | **0.04** | $COV=0.07$ | 0.05 | 0.18 | 0.06 | 0.02 | -1.29 | 0.22 |
| ARS | **[<0.001]** | [0.001] | [<0.001] | **[0.03]** | [0.05] | [0.05] | [0.001] | [<0.001] | [<0.001] | [-1.19] | [0.20] |
|  | **(<0.001, 0.05)** | (-0.04, 0.06) | (<0.001, 0.04) | **(0.01, 0.08)** | (0.01, 0.15) | (0.04, 0.07) | (<0.001, 0.38) | (<0.001, 0.15) | (<0.001, 0.05) | (-2.82, 0.23) | (-0.38, 0.84) |
| Male | $r_A=0.17$ | **$V_A=0.16$** | 0.05 | $r=0.67$ | **0.34** | 0.46 | 0.16 | 0.08 | 0.67 | -4.98 | 0.15 |
| ARS | $[0.18]$ | **[0.15]** | [<0.001] | $[0.78]$ | **[0.25]** | [0.46] | [0.12] | [0.08] | [0.40] | [-4.98] | [0.11] |
| | $(-0.57, 0.88)$ | **(0.02, 0.30)** | (<0.001, 0.15) | $(0.30, 0.96)$ | **(0.11, 0.64)** | (0.34, 0.58) | (0.03, 0.31) | (0.01, 0.15) | (0.01, 1.64) | (-7.94, -2.07) | (-1.18, 1.61) |
| Adult | 0.04 |  | 0.37 | 0.69 |  | 1 (fixed) | 0.02 | 0.005 | NA | -0.49 | 0.33 |
| annual | [<0.001] |  | [0.01] | [0.54] |  |  | [<0.001] | [<0.001] |  | [0.03] | [0.46] |
| survival | (<0.001, 0.16) |  | (<0.001, 1.44) | (0.20, 1.31) |  |  | (<0.001, 0.07) | (<0.001, 0.02) |  | (-4.81, 3.83) | (-1.48, 2.10) |

**Figure 4.**
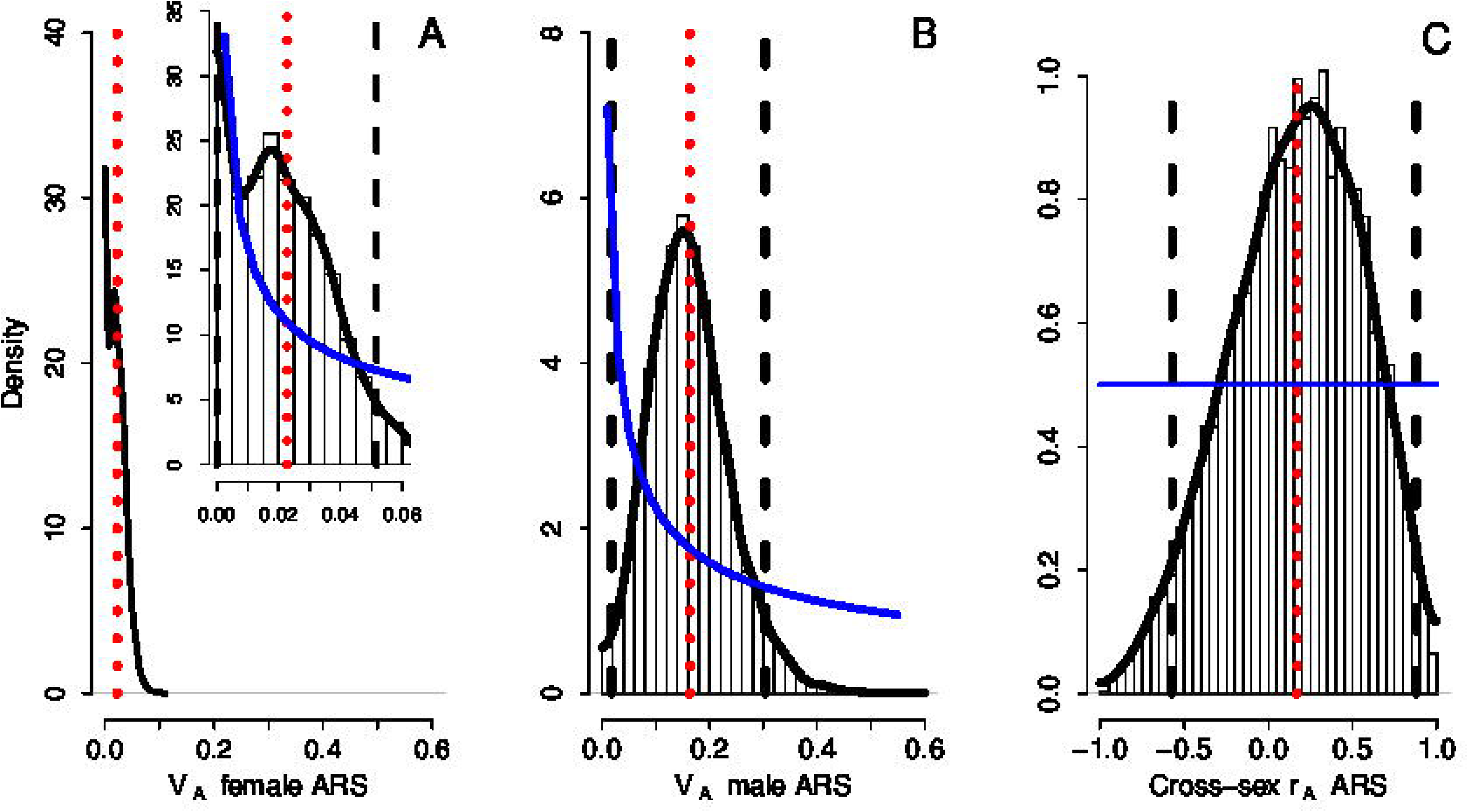
Marginal posterior MCMC samples (bars), kernel density estimation (solid black line), posterior mean (red dotted line), 95% credible interval limits (black dashed lines), and prior (solid blue line) for the additive genetic variances (V_A_) in (A) adult female annual reproductive success (ARS), (B) adult male ARS, and (C) the cross-sex additive genetic correlation (r_A_) in song sparrows. On A and B, x-axis scales are standardized to facilitate comparison, but the y-axis scales differ. The panel A inset shows the marginal posterior distribution for female ARS on a larger scale. In A and B, the priors are depicted over the range of each posterior distribution, but extend to substantial positive values.

In contrast, the posterior mode and mean for V_A_ in male ARS were substantially larger and the lower 95%CI limit did not converge towards zero (Table 3, Fig. 4B). The permanent individual variances were very small in both sexes, but the year and residual variances were substantial, especially for males (Table 3). Consequently, despite the marked difference in V_A_, the posterior means for 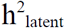 and 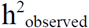 for ARS were similar in both sexes (~0.06–0.18), but I_A-observed_ was substantially greater in males than females (Table 3, Figs. S6, S7).

The posterior mode for the cross-sex additive genetic correlation (r_A_) in ARS was positive but small. Due to the small V_A_ in female ARS, the 95%CI was again wide and spanned zero, but did not converge towards either -1 or 1 (Table 3, Fig. 4C).

The posterior modes for the regressions of ARS on *IGG* were small in both sexes, with 95%CIs that overlapped zero (Table 3). The posterior modes for the regressions of ARS on *f* were negative in both sexes, although the 95%CI for females again overlapped zero (Table 3).

### ADULT SURVIVAL

For the focal 254 adult females and 331 adult males, the mean number of observations of annual survival (or mortality) was 2.1 (median 1, range 1–9, Fig. 5A) for females and 2.3 (median 2, range 1–9, Fig. 5B) for males, representing overall survival rates of 53.0% and 58.0% respectively.

**Figure 5.**
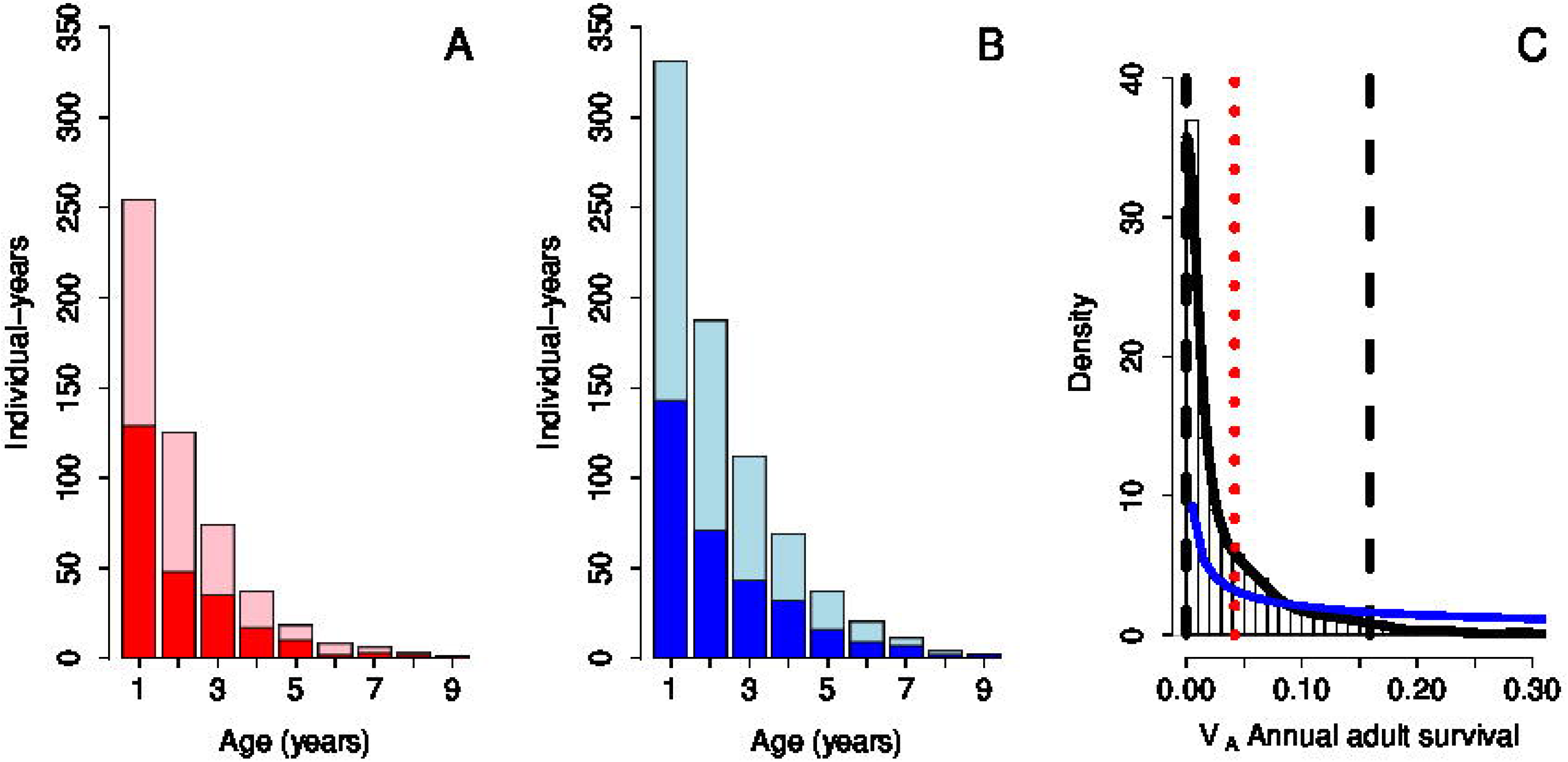
Phenotypic distributions of age-specific survival (or mortality) for adult (A) female and (B) male song sparrows, and (C) the marginal posterior distribution for additive genetic variance (V_A_) in annual adult survival. In A and B, dark and light shading indicate observations of mortality and survival respectively. In C, plot attributes are as for figures 2–4.

In the univariate QGGLMM, the posterior mode for V_A_ was effectively zero (Table 3, Fig. 5C). The posterior mean was slightly larger, but there was substantial posterior density close to zero compared to the prior distribution, and the lower 95%CI limit converged to zero (Table 3). Since there was also substantial year variance, the posterior modes for 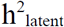 and 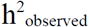 were very small (Table 3; Fig. S8.3). The posterior modes for the regressions of adult survival on *IGG* and *f* were also small, with 95%CIs that overlapped zero (Table 3). Analyses of adult longevity rather than annual survival yielded similar conclusions (Appendix S8).

## Discussion

### ADDITIVE GENETIC VARIANCE AND CORRELATION IN SEX-SPECIFIC FITNESS

The sex-specific additive genetic variances (V_A_) in fitness, and the cross-sex genetic correlation (r_A_), are key parameters that determine the rate of fitness evolution and shape evolutionary responses to natural and sexual selection (Burt 1995; Brommer et al. 2007; Kirkpatrick 2009; Shaw and Shaw 2014). They also underlie the potential for evolutionary sexual conflict, which might constrain evolution yet help maintain overall V_A_ in fitness (Lande 1980; Chippindale et al. 2001; Kruuk et al. 2008; Bonduriansky and Chenoweth 2009; Long et al. 2012). However, these key parameters have rarely been estimated in wild populations, particularly using theoretically appropriate measures of fitness while accommodating non-Gaussian phenotypic distributions and accounting for genetic effects of immigration and inbreeding (Kruuk et al. 2008; Kirkpatrick 2009; Shaw and Etterson 2012; Gomulkiewicz and Shaw 2013; Shaw and Shaw 2014).

Our analyses of comprehensive fitness data from free-living song sparrows estimated nonzero latent-scale V_A_s and heritabilities for fitness, of similar magnitudes, in both sexes. Such estimates do not concur with basic theoretical predictions that V_A_ in fitness will be negligible at equilibrium (Charlesworth 1987), which has been interpreted to commonly apply (Shaw and Shaw 2014; Walling et al. 2014). Instead, they support the view that substantial V_A_ in fitness can be readily generated and/or maintained and imply that this population is not at an evolutionary equilibrium (Houle 1992; Kirkpatrick 2009; Zhang 2012; Shaw and Shaw 2014). Further, our estimate of a moderate positive cross-sex r_A_ for fitness implies that some V_A_ is shared between the sexes, potentially facilitating an increase in population mean fitness (Lande 1980). However, the cross-sex r_A_ in fitness was detectably less than one, implying that some sexually antagonistic genetic variation does exist, potentially facilitating the maintenance of overall V_A_.

The few available estimates of sex-specific V_A_ in fitness in wild populations cannot readily be quantitatively compared because different studies used different fitness metrics, analytical methods and estimation scales, with different degrees of paternity error and missing data. However, qualitatively concordant with our results, V_A_ for fitness was estimated to be non-zero and similar in both sexes in collared flycatchers (*Ficedula albicollis*, Merilä & Sheldon 2000; Brommer et al. 2007) and Swedish humans (Zietsch et al. 2014). Conversely, V_A_ was estimated to be zero or very small in both sexes in great tits (*Parus major*, McCleery et al. 2004), bighorn sheep (*Ovis canadensis*, Coltman et al. 2005), North American red squirrels (*Tamiasciurus hudsonicus*, McFarlane et al. 2014), and savannah sparrows (*Passerculus sandwichensis*, Wheelwright et al. 2014); zero in females but more substantial in males in red deer (*Cervus elaphus*, Kruuk et al. 2000, but see Foerster et al. 2007) and Austrian humans (Gavrus-Ion et al. 2017); yet zero in males but more substantial in females in red-billed gulls (*Larus novaehollandiae*, Teplitsky et al. 2009) and pre-industrial Finnish humans (Pettay et al. 2005, Appendix S1).

Meanwhile, our estimate of a moderate positive cross-sex r_A_ for fitness differs from the substantial negative values previously estimated in wild populations (Foerster et al. 2007; Brommer et al. 2007; McFarlane et al. 2014; Appendix S1), and from the small or slightly negative values estimated in laboratory populations (Chippindale et al. 2001; Delcourt et al. 2009; Innocenti and Morrow 2010; Collet et al. 2016). Yet, cross-sex r_A_s can change substantially when (laboratory) populations experience novel environments (Delcourt et al. 2009; Punzalan et al. 2014; Collet et al. 2016), migration load (Long et al. 2012), or inbreeding (Duffy et al. 2014). Positive estimates, such as ours, might indicate populations where both sexes are displaced from their fitness peak, and consequently experience congruent directional selection (Long et al. 2012; Duffy et al. 2014; Punzalan et al. 2014). Overall, further rigorous and standardized estimates of V_A_ and r_A_ in sex-specific fitness from wild populations experiencing different ecological circumstances are clearly required to discern general patterns and evolutionary implications.

### ADDITIVE GENETIC VARIANCES AND CORRELATIONS IN FITNESS COMPONENTS

Values of V_A_ in sex-specific fitness, and the cross-sex r_A_, must ultimately result from V_A_s and crosssex and within-sex r_A_s in underlying sex-specific fitness components. Quantifying such parameters can consequently help identify mechanisms that maintain V_A_ in fitness, and identify sources of sexual conflict (Walling et al. 2014). Juvenile survival to maturity constitutes one primary fitness component; indeed, 96% of observed song sparrow fitness values of zero represent individuals that did not (locally) survive to adulthood, and such patterns are likely commonplace (Blomquist 2010; Wagenius et al. 2010; Gomulkiewicz and Shaw 2013). We estimated moderate V_A_ in juvenile survival, concurring with previous evidence that V_A_ is moderate and similar in female and male song sparrows with a substantial positive cross-sex r_A_ (Reid and Sardell 2012, Appendix S7). However, for adult LRS, which constitutes the remaining primary fitness component, there was a striking difference between the sexes: V_A_ for male LRS was substantial and clearly exceeded zero, while V_A_ for female LRS was very small. This implies that there is opportunity for relatively rapid evolutionary change in male LRS and genetically correlated traits, but little such opportunity regarding female LRS.

The small V_A_ estimate for female LRS impedes precise estimation of the cross-sex r_A_ in LRS, and indeed renders such estimation somewhat redundant (since r_A_ is undefined given zero V_A_ in one or both sexes). Nevertheless, the posterior mode was small, and if anything slightly negative, further suggesting that additive genetic effects on adult LRS are largely independent in females and males. Together, our results imply that the moderate positive cross-sex r_A_ in fitness is primarily driven by the positive cross-sex r_A_ in juvenile survival. Consequently, the cross-sex expression of additive genetic effects on juvenile survival ameliorates potentially sexually antagonistic genetic variation in overall fitness resulting from sex-specific expression of adult LRS. These patterns are reminiscent of those observed in *Drosophila melanogaster*, where a positive cross-sex r_A_ in juvenile survival initially combined with a negative cross-sex r_A_ in adult reproductive success to generate a weak overall cross-sex r_A_ for fitness (Chippindale et al. 2001), but where the cross-sex r_A_ in adult reproductive success was no longer detectably different from zero after further generations of laboratory adaptation (Collet et al. 2016).

Further decomposition of adult LRS in song sparrows revealed little or no V_A_ in adult annual survival, and identified ARS as the primary source of V_A_ in male LRS. The substantial difference in V_A_ in ARS, and hence LRS, between males and females likely reflects the population’s ecology and mating system. Due to the typically male-biased adult sex-ratio and frequent extra-pair paternity, males accumulate ARS by securing a territory and a social mate, defending within-pair paternity and accruing extra-pair paternity (Sardell et al. 2010; Lebigre et al. 2012; Reid et al. 2011, 2014a,b). In contrast, females accumulate ARS through their own fecundity. Consequently, while components of ARS such as within-pair paternity can be conceptualized as ‘emergent’ traits of pairs rather than individuals (Reid et al. 2014a), males and females are likely to differ substantially in the suite of physiological and behavioral traits that generate high ARS, and hence in underlying genetic effects. Previous analyses revealed non-zero V_A_ in annual male extra-pair reproductive success and a positive r_A_ with within-pair paternity success per brood (Reid et al. 2014b), but a negative r_A_ between net paternity success and juvenile survival (Reid and Sardell 2012). Together, these positive and negative correlations, alongside among-year variation in adult sex-ratio and hence the social context in which male reproductive success is expressed, could help maintain substantial V_A_ in male ARS (and hence LRS).

### GENETIC EFFECTS OF IMMIGRATION

Immigration, and resulting gene flow, is one primary mechanism that can maintain V_A_ in fitness and associated evolutionary potential in any focal population, and can also rapidly increase mean fitness by alleviating inbreeding depression. However the overall genetic effects of immigration, and the evolutionary consequences, depend on genetic properties of naturally-occurring immigrants compared to existing natives (Ingvarsson and Whitlock 2000; Tallmon et al. 2004; Edelaar and Bolnick 2012; Carlson et al. 2014). We utilized the multi-generation song sparrow pedigree, that links all Mandarte-hatched individuals to their immigrant and ‘founder’ ancestors and hence describes the expected introgression of immigrants’ alleles, to directly estimate the relative mean additive genetic values for local fitness of the defined immigrant and founder genetic groups.

Unlike analyses that aim to discern demographic and evolutionary consequences of dispersal by directly comparing observed phenotypes of immigrants (or dispersers) and residents (e.g. Marr et al. 2002; Pasinelli et al. 2004; Nosil et al. 2005; Pärn et al. 2009; Bonte et al. 2012), our analyses do not utilize immigrants’ own phenotypes and consequently cannot be confounded by environmental effects of dispersal on those phenotypes. Our analyses showed that immigrant song sparrows carry alleles that, when expressed in subsequent Mandarte-hatched generations, have negative additive effects on local fitness in both sexes.

Such negative effects of immigrant’s alleles could stem from three main processes. First, there could be divergent selection among song sparrow demes and resulting local adaptation. Immigrants to Mandarte might consequently not be locally adapted and hence have low mean additive genetic value for local fitness, as assumed by classical migration load models. Second, dispersal could be non-random, such that individuals that immigrate into Mandarte have low additive genetic value for local fitness even in the absence of any local adaptation. Third, low additive genetic value for fitness measured on Mandarte could reflect V_A_ in dispersal, such that offspring of immigrants are more likely to emigrate and hence have zero local fitness (e.g. Doligez and Pärt 2008). These three processes, which are not mutually exclusive, are not distinguished by our current analyses.

However, immigrants’ low additive genetic value for fitness resulted primarily from low additive genetic value for local juvenile survival rather than subsequent adult LRS, and therefore reflects some combination of effects on early-life mortality and/or emigration. To indicate biological effect sizes, the estimated latent-scale effect of *β*_IGG_ = -2.6 (Table 2) implies a decrease in local juvenile survival probability of approximately 0.04 given an increase in individual *IGG* coefficient of 0.1 spanning the current mean of ~0.5, which is not trivial. In general, such reduced local survival of immigrants’ descendants would reduce the effective rate of gene flow below that expected given the observed immigration rate (Garant et al. 2007).

However, our analyses also demonstrate strong inbreeding depression in fitness in both sexes, resulting from inbreeding depression in juvenile survival and adult LRS and ARS (as previously documented, Keller 1998; Reid et al. 2014c; Nietlisbach et al. 2017). Such inbreeding depression reflects covariance between individual fitness and *f*, where the underlying variance in *f* stems substantially from immigrant-native outcrossing; resulting F1 offspring are defined as outbred and have relatively high fitness (Keller 1998; Marr et al. 2002; Reid et al. 2006, 2014c; Wolak and Reid 2016). The estimated latent-scale effect size of *β*_f_ = -9.3 (Table 2) implies an increase in juvenile survival probability of approximately 0.25 for outbred offspring (*f*=0) compared to inbred offspring with *f*=0.1 (see also Keller 1998; Reid et al. 2014c). This effect could cause rapid initial introgression of immigrants’ alleles, and hence increase the short-term effective rate of gene flow (e.g. Ingvarsson and Whitlock 2000; Garant et al. 2007; Hedrick et al. 2014). Indeed, the mean *IGG* coefficient of ~0.5, calculated across the focal 2821 fitness-phenotyped individuals, implies that an average Mandarte-hatched song sparrow inherited half its genome from immigrant ancestors despite the relatively small number of contributing immigrants (n=26) and that only 195 (7%) of the phenotyped individuals were direct F1 offspring of immigrant-native pairings.

However, once immigrants’ descendants start to inbreed, as is inevitable for initially high-fitness lineages in small populations (Reid et al. 2006; Bijlsma et al. 2010; Hedrick et al. 2014), increased expression of recessive alleles with detrimental effects on local fitness would occur. This process would exacerbate the decrease in fitness that is expected following recombination in F2 and subsequent generations and resulting outbreeding depression (Frankham 2016), as documented in song sparrows (Marr et al. 2002). The combination of heterosis that exacerbates initial introgression and low overall additive genetic value for fitness could potentially generate substantial migration load; almost all population members might be pulled below the fitness peak, substantially decreasing population mean fitness but potentially generating a positive overall cross-sex r_A_ for fitness (as observed in song sparrows) and alleviating sexual conflict (Long et al. 2012; Duffy et al. 2014; Punzalan et al. 2014). Such multi-generational dynamics of immigrants’ alleles can, in future, be explicitly quantified using pedigree and genomic data (from song sparrows and other systems), and through theory that simultaneously considers heterosis and migration load (e.g. Lopez et al. 2009). Meanwhile, our analyses demonstrate that structured quantitative genetic analyses can explicitly estimate V_A_ in fitness alongside multiple genetic consequences of immigration in wild populations, and thereby inform understanding of the contributions of gene flow to the magnitude and maintenance of overall V_A_ in fitness and resulting evolutionary dynamics.

## AUTHOR CONTRIBUTIONS

MEW and JMR jointly conceived and designed the analyses and wrote the initial draft of the manuscript. MEW conducted the analyses, with contributions from JMR. PA ensured the long-term field data collection. PN and LFK conducted the genotyping. All authors contributed to editing and writing of the final draft.

## ACKNOWLEDGMENTS

We thank the Tsawout and Tseycum First Nation bands for access to Mandarte, Pirmin Nietlisbach, and everyone who contributed to the long-term data collection. We thank the European Research Council for funding and the University of Aberdeen for generous access to the Maxwell High Performance Computing cluster. Pierre de Villemereuil, Michael B. Morrissey, and Jarrod D. Hadfield provided enlightening discussions during manuscript preparation.

## Supporting Information

Additional supporting information may be found in the online version of this article at the publisher’s website:

**Appendix S1**. Literature summary

**Appendix S2**. Overall approach and data specifications

**Appendix S3**. Pedigree structure and genetic groups

**Appendix S4**. Details of model specification and implementation

**Appendix S5**. Additional details of results

**Appendix S6**. Prior sensitivity analysis

**Appendix S7**. Sex-specific juvenile survival

**Appendix S8**. Additional analyses of adult survival

## LITERATURE CITED

Arnold, S. J., and M. J. Wade. 1984. On the measurement of natural and sexual selection: Applications. Evolution 38: 720–734.

Barton, N. H., and P. D. Keightley. 2002. Understanding quantitative genetic variation. Nat. Rev. Gen. 3: 11–21.

Bell, G. 2013. Evolutionary rescue and the limits of adaptation. Philos. Trans. R. Soc. B 368: 20120080.

Bijlsma, R., M. D. D. Westerhof, L. P. Roekx, and I. Pen. 2010. Dynamics of genetic rescue in inbred Drosophila melanogaster population. Cons. Genet. 11: 449–462.

Blomquist, G.E. 2010. Heritability of fitness in female macaques. Evol. Ecol. 24:657–669.

Bonduriansky, R., and S. F. Chenoweth. 2009. Intralocus sexual conflict. Trends Ecol. Evol. 24: 280–288.

Bonte, D. et al. 2012. Costs of dispersal. Biol. Rev. 87: 290–312.

Brommer, J. E. 2000. The evolution of fitness in life-history theory. Biol. Rev. 75: 377–404.

Brommer, J. E., M. Kirkpatrick, A. Qvarnström, and L. Gustafsson. 2007. The intersexual genetic correlation for lifetime fitness in the wild and its implications for sexual selection. PLoS One 2:e744.

Burt, A. 1995. The evolution of fitness. Evolution, 49: 1–8.

Carlson, S. M., C. J. Cunningham, and P. A. H. Westley. 2014. Evolutionary rescue in a changing world. Trends. Ecol. Evol. 29: 521–530.

Charlesworth, B. 1987. The heritability of fitness. Pp. 21–40 *in* J. W. Bradbury and M. B. Andersson, eds. Sexual Selection: Testing the Alternatives. Wiley, Chichester.

Charmantier, A., D. Garant, and L. E. B. Kruuk (eds). 2014. Quantitative Genetics in the Wild. Oxford University Press, Oxford.

Chippindale, A. K., J. R. Gibson, and W. R. Rice. 2001. Negative genetic correlation for adult fitness between sexes reveals ontogenetic conflict in *Drosophila*. Proc. Natl. Acad. Sci. U. S. A. 98: 1671–1675.

Collet, J. M., S. Fuentes, J. Hesketh, M. S. Hill, P. Innocenti, E. H. Morrow, K. Fowler, and M. Reuter. 2016. Rapid evolution of the intersexual genetic correlation for fitness in *Drosophila melanogaster*. Evolution 70: 781–795.

Coltman, D. W., P. O’Donoghue, J. T. Hogg, and M. Festa-Bianchet. 2005. Selection and genetic (co)variance in bighorn sheep. Evolution 59: 1372–1382.

Crow, J. F., and M. Kimura. 1970. Introduction to Population Genetics Theory. Harper and Row, New York.

de Villemereuil, P., H. Schielzeth, S. Nakagawa, and M. Morrissey. 2016. General methods for evolutionary quantitative genetic inference from generalized mixed models. Genetics 204: 1281–1294.

Delcourt, M., M. W. Blows, and H. D. Rundle. 2009. Sexually antagonistic genetic variance for fitness in an ancestral and a novel environment. Proc. R. Soc. B 276: 2009–14.

Doligez, B., and T. Pärt. 2008. Estimating fitness consequences of dispersal: a road to ‘know-where’? Non-random dispersal and the underestimation of dispersers’ fitness. J. Anim. Ecol. 77: 1199–1211.

Duffy, E., R. Joag, J. Radwan, N. Wedell, and D. J. Hosken. 2014. Inbreeding alters intersexual fitness correlations in *Drosophila simulans*. Ecol. and Evol. 4: 3330–3338.

Edelaar, P., and D. I. Bolnick. 2012. Non-random gene flow: an underappreciated force in evolution and ecology. Trends Ecol. Evol. 27: 659–665.

Ellegren, H., and B. C. Sheldon. 2008. Genetic basis of fitness differences in natural populations. Nature 452: 169–175.

Falconer, D. S. 1989. Introduction to Quantitative Genetics. 3rd ed. John Wiley & Sons, Inc., New York.

Firth, J. A., J. D. Hadfield, A. W. Santure, J. Slate, and B. C. Sheldon. 2015. The influences of nonrandom extra-pair paternity on heritability estimates derived from wild pedigrees. Evolution 69: 1336–1344.

Fisher, R. A. 1930. The Genetical Theory of Natural Selection. Clarendon Press, Oxford.

Flint, J., and T. F. C. Mackay. 2009. Genetic architecture of quantitative traits in mice, flies, and humans. Genome Res. 19: 723–733.

Foerster, K., T. Coulson, B. C. Sheldon, J. M. Pemberton, T. H. Clutton-Brock, and L. E. B. Kruuk. 2007. Sexually antagonistic genetic variation for fitness in red deer. Nature 447: 1107–1111.

Frankham, R. 2016. Genetic rescue benefits persist to at least the F3 generation, based on metaanalysis. Biol. Cons. 195, 33–36

Freeman-Gallant, C. R., N. T. Wheelwright, K. E. Meiklejohn, S. L. States, and S. V Sollecito. 2005. Little effect of extrapair paternity on the opportunity for sexual selection in Savannah sparrows (*Passerculus sandwichensis*). Evolution 59: 422–430.

Garant, D., S. E. Forde, and A. P. Hendry. 2007. The multifarious effects of dispersal and gene flow on contemporary adaptation. Func. Ecol. 21: 434–443.

Garcia-Gonzalez, F., L. W. Simmons, J. L. Tompkins, J. S. Kotiaho, and J. P. Evans. 2012. Comparing evolvabilities: common errors surrounding the calculation and use of coefficients of additive genetic variation. Evolution 60, 2341–2349.

Gardner, M. P., K. Fowler, N. H. Barton, and L. Partridge. 2005. Genetic variation for total fitness in *Drosophila melanogaster:* Complex yet replicable patterns. Genetics 169: 1553–1571.

Gavrus-Ion, A., T. Sjøvold, M. Hernández, R. González-José, M. E. Esteban Torné, N. Martínez-Abadías, and M. Esparza. 2017. Measuring fitness heritability: Life history traits versus morphological traits in humans. Am. J. Phys. Anthropol. 164: 321–330.

Gelman, A. 2006. Prior distributions for variance parameters in hierarchical models. Bayesian Anal. 1: 515–533.

Gomulkiewicz, R., and R. G. Shaw. 2013. Evolutionary rescue beyond the models. Philos. Trans. R. Soc. B 368: 20120093.

Hadfield, J. D. 2008. Estimating evolutionary parameters when viability selection is operating. Proc. R. Soc. B 275: 723–734.

Hadfield, J. D. 2010. MCMC methods for multi-response generalized linear mixed models: the MCMCglmm R package. J. Stat. Soft. 33: 1–22.

Hadfield, J. D., E. A. Heap, F. Bayer, E. A. Mittell, and N. M. A. Crouch. 2013. Disentangling genetic and prenatal sources of familial resemblance across ontogeny in a wild passerine. Evolution 67: 2701–2713.

Hedrick, P. W., R. O. Peterson, L. M. Vucetich, J. R. Adams, and J. A. Vucetich. 2014. Genetic rescue in Isle Royale wolves: genetic analysis and the collapse of the population. Cons. Genet. 15: 1111–1121.

Henderson, C. R. 1973. Sire evaluation and genetic trends. J. Anim. Sci. 1973: 10–41.

Hill, W. G. 2012. Quantitative genetics in the genomics era. Curr. Genomics 13: 196–206.

Houle, D. 1992. Comparing evolvability and variability of quantitative traits. Genetics 130: 195–204.

Innocenti, P., and E. H. Morrow. 2010. The sexually antagonistic genes of *Drosophila melanogaster*. PLoS Biol. 8:e1000335.

Ingvarsson, P. K., and M. C. Whitlock. 2000. Heterosis increases the effective migration rate. Proc. R. Soc. B. 267: 1321–1326.

Johnston, S. E., J. Gratten, C. Berenos, J. G. Pilkington, T. H. Clutton-brock, J. M. Pemberton, and J. Slate. 2013. Life history trade-offs at a single locus maintain sexually selected genetic variation. Nature 502: 93–95.

Keller, L. F. 1998. Inbreeding and its fitness effects in an insular population of song sparrows (*Melospiza melodia*). Evolution 52: 240–250.

Keller, L. F., J. M. Reid, and P. Arcese. 2008. Testing evolutionary models of senescence in a natural population: age and inbreeding effects on fitness components in song sparrows. Proc. R. Soc. B 275: 597–604.

Kirkpatrick, M. 2009. Patterns of quantitative genetic variation in multiple dimensions. Genetica 136: 271–284.

Kruuk, L. E. B., T. H. Clutton-Brock, J. Slate, J. M. Pemberton, S. Brotherstone, and F. E. Guinness. 2000. Heritability of fitness in a wild mammal population. Proc. Natl. Acad. Sci. 97: 698–703.

Kruuk, L. E. B., A. Charmantier, and D. Garant. 2014. The study of quantitative genetics in wild populations. Pp. 1–15 in A. Charmantier, D. Garant, and L. E. B. Kruuk, eds. Quantitative Genetics in the Wild. Oxford University Press, Oxford.

Kruuk, L. E. B., and J. D. Hadfield. 2007. How to separate genetic and environmental causes of similarity between relatives. J. Evol. Biol. 20: 1890–1903.

Kruuk, L. E. B. 2004. Estimating genetic parameters in natural populations using the “animal model.” Philos. Trans. R. Soc. B 359: 873–890.

Kruuk, L. E. B., J. Slate, and A. J. Wilson. 2008. New answers for old questions: The evolutionary quantitative genetics of wild animal populations. Annu. Rev. Ecol. Evol. Syst. 39: 525–548.

Lande, R. 1980. Sexual dimorphism, sexual selection, and adaptation in polygenic characters. Evolution 34: 292–305.

Lande, R. 1982. A quantitative genetic theory of life history evolution. Ecology 63: 607–615.

Lebigre, C., P. Arcese, R. J. Sardell, L. F. Keller, and J. M. Reid. 2012. Extra-pair paternity and the variance in male fitness in song sparrows (*Melospiza melodia*). Evolution 66: 3111–3129.

Lenormand, T. 2002. Gene flow and the limits to natural selection. Trends Ecol. Evol. 17: 183–189.

Lewontin, R. C. 1974. Genetic Basis of Evolutionary Change. Columbia University Press, New York.

Long, T. A. F., A. F. Agrawal, and L. Rowe. 2012. The effect of sexual selection on offspring fitness depends on the nature of genetic variation. Cur. Biol. 22: 204–208.

Lopez, S., F. Rousset, F. H. Shaw, R. G. Shaw, and O. Ronce. 2009. Joint effects of inbreeding and local adaptation on the evolution of genetic load after fragmentation. Cons. Biol. 23: 1618–1627.

Lynch, M., and B. Walsh. 1998. Genetics and analysis of quantitative traits. Sinauer, Sunderland, USA.

Marr, A. B., L. F. Keller, and P. Arcese. 2002. Heterosis and outbreeding depression in descendants of natural immigrants to an inbred population of song sparrows (Melospiza melodia). Evolution 56: 131–142.

McCleery, R. H., R. A. Pettifor, P. Armbruster, K. Meyer, B. C. Sheldon, and C. M. Perrins. 2004. Components of variance underlying fitness in a natural population of the great tit *Parus major*. Am. Nat. 164:E62–E72.

McFarlane, S. E., J. C. Gorrell, D. W. Coltman, M. M. Humphries, S. Boutin, and A. G. McAdam. 2014. Very low levels of direct additive genetic variance in fitness and fitness components in a red squirrel population. Ecol. & Evol. 4: 1729–1738

Merilä, J., and B. C. Sheldon. 1999. Genetic architecture of fitness and nonfitness traits: empirical patterns and development of ideas. Heredity 83: 103–109.

Merilä, J., and B. C. Sheldon. 2000. Lifetime reproductive success and heritability in nature. Am. Nat. 155: 301–310.

Metcalf, C. J. E., and S. Pavard. 2007. Why evolutionary biologists should be demographers. Trends Ecol. Evol. 22: 205–212.

Milot, E., F. M. Mayer, D. H. Nussey, M. Boisvert, F. Pelletier, and D. Réale. 2011. Evidence for evolution in response to natural selection in a contemporary human population. Proc. Natl. Acad. Sci. 108: 17040–45

Nietlisbach, P., G. Camenisch, T. Bucher, J. Slate, L. F. Keller, and E. Postma. 2015. A microsatellite-based linkage map for song sparrows (*Melospiza melodia*). Mol. Ecol. Res. 15: 1486–1496.

Nietlisbach, P., L. F. Keller, G. Camenisch, F. Guillaume, P. Arcese, J. M. Reid, and E. Postma. 2017. Pedigree-based inbreeding coefficient explains more variation in fitness than heterozygosity at 160 microsatellites in a wild bird population. Proc. R. Soc. B 284: 20162763.

Nosil, P., T. H. Vines, and D. J. Funk. 2005. Perspective: Reproductive isolation caused by natural selection against immigrants from divergent habitats. Evolution 59: 705–719.

Orr, H. A. 2009. Fitness and its role in evolutionary genetics. Nat. Rev. Genet. 10: 531–539.

Pärn, H., H. Jensen, T. H. Ringsby, and B.-E. Sæther. 2009. Sex-specific fitness correlates of dispersal in a house sparrow metapopulation. J. Anim. Ecol. 78: 1216–1225.

Pasinelli, G., K. Schiegg, and J. R. Walters. 2004. Genetic and environmental influences on natal dispersal distance in a resident bird species. Am. Nat. 164: 660–669.

Pettay, J. E., Kruuk, L. E. B., J. Jokela, and V. Lummaa. 2005. Heritability and genetic constraints of life-history evolution in preindustrial humans. Proc. Natl. Acad. Sci. 102: 2838–2843.

Poissant, J., A. J. Wilson, and D. W. Coltman. 2010. Sex-specific genetic variance and the evolution of sexual dimorphism: a systematic review of cross-sex genetic correlations. Evolution 64: 97–107.

Postma, E., F. Heinrich, U. Koller, R. J. Sardell, J. M. Reid, P. Arcese, and L. F. Keller. 2011. Disentangling the effect of genes, the environment and chance on sex ratio variation in a wild bird population. Proc. R. Soc. B 278: 2996–3002.

Price, G. R. 1970. Selection and covariance. Nature 227: 520–521.

Punzalan, D., M. Delcourt, and H. D. Rundle. 2014. Comparing the intersex genetic correlation for fitness across novel environments in the fruit fly, *Drosophila serrata*. Heredity 112: 143–148.

R Core Team. 2015. R: A language and environment for statistical computing. R Foundation for Statistical Computing, Vienna, Austria.

Reid, J. M., P. Arcese, and L. F. Keller. 2006. Intrinsic parent-offspring correlation in inbreeding level in a population open to immigration. Am. Nat. 168: 1–13.

Reid, J. M., P. Arcese, L. F. Keller, and S. Losdat. 2014a. Female and male genetic effects on offspring paternity: additive genetic (co)variances in female extra-pair reproduction and male paternity success in song sparrows (*Melospiza melodia*). Evolution 68: 2357–2370.

Reid, J. M., P. Arcese, and S. Losdat. 2014b. Genetic covariance between components of male reproductive success: within-pair versus extra-pair paternity in song sparrows. J. Evol. Biol. 27: 2046–2056.

Reid, J. M., P. Arcese, R. J. Sardell, and L. F. Keller. 2011. Additive genetic variance, heritability and inbreeding depression in male extra-pair reproductive success. Am. Nat. 177: 177–187.

Reid, J. M., and L. F. Keller. 2010. Correlated inbreeding among relatives: occurrence, magnitude, and implications. Evolution 64: 973–985.

Reid, J. M., L. F. Keller, A. B. Marr, P. Nietlisbach, R. J. Sardell, and P. Arcese. 2014c. Pedigree error due to extra-pair reproduction substantially biases estimates of inbreeding depression. Evolution 68: 802–815.

Reid, J. M., and R. J. Sardell. 2012. Indirect selection on female extra-pair reproduction? Comparing the additive genetic value of maternal half-sib extra-pair and within-pair offspring. Proc. R. Soc. B 279: 1700–1708.

Robertson, A. 1966. A mathematical model of the culling process in dairy cattle. Anim. Prod. 8: 95–108.

Rose, M. R. 1982. Antagonistic pleiotropy, dominance, and genetic variation. Heredity 48: 63–78.

Sæther, B.-E., and S. Engen. 2015. The concept of fitness in fluctuating environments. Trends Ecol. Evol. 30: 273–281.

Sardell, R. J., L. F. Keller, P. Arcese, T. Bucher, and J. M. Reid. 2010. Comprehensive paternity assignment: genotype, spatial location and social status in song sparrows, *Melospiza melodia*. Mol. Ecol. 19: 4352–4364.

Shaw, R. G. 1987. Maximum-likelihood approaches applied to quantitative genetics of natural populations. Evolution 41: 812–826.

Shaw, R. G., and J. R. Etterson. 2012. Rapid climate change and the rate of adaptation: insight from experimental quantitative genetics. New Phytol. 195: 752–765.

Shaw, R. G., and F. H. Shaw. 2014. Quantitative genetic study of the adaptive process. Heredity 112: 13–20.

Smith, J. N. M., L. F. Keller, A. B. Marr, and P. Arcese (eds). 2006. Conservation and Biology of Small Populations: The Song Sparrows of Mandarte Island. Oxford University Press, Oxford.

Stinchcombe, J. R. 2014. Cross-pollination of plants and animals: wild quantitative genetics and plant evolutionary genetics. Pp. 128–146 *in* A. Charmantier, D. Garant, and L. E. B. Kruuk, eds. Quantitative Genetics in the Wild. Oxford University Press, Oxford, UK.

Tallmon, D. A., G. Luikart, and R. S. Waples. 2004. The alluring simplicity and complex reality of genetic rescue. Trends Ecol. Evol. 19: 489–496.

Teplitsky, C., J. A. Mills, J. W. Yarrall, and J. Merilä. 2009. Heritability of fitness components in a wild bird population. Evolution 63: 716–726.

Trask, A. E., E. M. Bignal, D. I. McCracken, P. Monaghan, S. B. Piertney, and J. M. Reid. 2016. Evidence of the phenotypic expression of a lethal recessive allele under inbreeding in a wild population of conservation concern. J. Anim. Ecol. 85: 879–891.

Travisano, M., and R. G. Shaw. 2013. Lost in the map. Evolution 67: 305–314.

Wagenius, S., H. H. Hangelbroek, C. E. Ridley, and R. G. Shaw. 2010. Biparental inbreeding and interremnant mating in a perennial prairie plant: fitness consequences for progeny in their first eight years. Evolution 64: 761–771.

Walling, C. A., M. B. Morrissey, K. Foerster, T. H. Clutton-Brock, J. M. Pemberton, and L. E. B. Kruuk. 2014. A multivariate analysis of genetic constraints to life history evolution in a wild population of red deer. Genetics 198: 1735–1749.

Walsh, B., and M. W. Blows. 2009. Abundant genetic variation + strong selection = multivariate genetic constraints: a geometric view of adaptation. Annu. Rev. Ecol. Evol. Syst. 40: 41–59.

Wheelwright, N. T., L. F. Keller, and E. Postma. 2014. The effect of trait type and strength of selection on heritability and evolvability in an island bird population. Evolution 68: 3325–3336.

Wilson, S., D. R. Norris, A. G. Wilson, and P. Arcese. 2007. Breeding experience and population density affect the ability of a songbird to respond to future climate variation. Proc. R. Soc. B 274: 2539–2545.

Wolak, M. E. 2012. nadiv: an R package to create relatedness matrices for estimating non-additive genetic variances in animal models. Methods Ecol. Evol. 3: 792–796.

Wolak, M. E., and L. F. Keller. 2014. Dominance genetic variance and inbreeding in natural populations. Pp. 104–127 *in* A. Charmantier, D. Garant, and L. E. B. Kruuk, eds. Quantitative Genetics in the Wild. Oxford University Press, Oxford.

Wolak, M. E., and J. M. Reid. 2016. Is pairing with a relative heritable? Estimating female and male genetic contributions to the degree of biparental inbreeding in song sparrows (*Melospiza melodia*). Am. Nat. 187: 736–752.

Wolak, M. E., and J. M. Reid. 2017. Accounting for genetic differences among unknown parents in microevolutionary studies: how to include genetic groups in quantitative genetic animal models. J. Anim. Ecol. 86: 7–20.

Wolak, M. E., D. A. Roff, and D. J. Fairbairn. 2015. Are we underestimating the genetic variances of dimorphic traits? Ecol. Evol. 5: 590–597.

Wolf, J. B., and M. J. Wade. 2001. On the assignment of fitness to parents and offspring: whose fitness is it and when does it matter? J. Evol. Biol. 14: 347–356.

Zhang, X.-S. 2012. Fisher’s geometrical model of fitness landscape and variance in fitness within a changing environment. Evolution 66: 2350–2368.

Zietsch, B. P., R. Kuja-Halkola, H. Walum, and K. J. H. Verweij. 2014. Perfect genetic correlation between number of offspring and grandoffspring in an industrialized human population. Proc. Natl. Acad. Sci. 111: 1032–1036.

